# Beyond viral detection: multitrophic effects of covert infection with an RNA virus in medfly

**DOI:** 10.1101/2022.11.25.517915

**Authors:** Luis Hernández-Pelegrín, Ricardo García-Martínez, Elena Llácer, Lorena Nieves, Ángel Llopis-Giménez, Marta Catalá-Oltra, Óscar Dembilio, Meritxell Pérez-Hedo, Alberto Urbaneja, Vera I.D. Ros, Francisco Beitia, Salvador Herrero

## Abstract

**Background:** With the advent of high-throughput sequencing, large sets of insect-infecting RNA viruses producing apparent asymptomatic infections are being discovered. In the Mediterranean fruit fly (medfly) *Ceratitis capitata*, an agricultural key pest of a wide range of fruits, up to 13 different RNA viruses have been described. Recent analysis demonstrated a wide distribution of these viruses in different medfly strains collected worldwide, but little is known about the interactions between those viruses and the medfly host. Previous studies suggested that a higher abundance of Ceratitis capitata nora virus (CcaNV) decreased medfly developmental time. Here, we investigated the effect of CcaNV on a broad range of parameters related to host fitness and its interaction with other trophic levels.

**Results:** CcaNV purified from a naturally infected medfly strain was used to infect CcaNV-free flies orally and subsequently monitor pupal weight, adult emergence, flying ability and longevity. Our results revealed detrimental effects associated with a CcaNV infection in the medfly, in terms of reduced pupal weight and reduced adult longevity. Moreover, we tested the influence of a CcaNV infection in medflies on the parasitism performance of *Aganaspis daci*, a medfly endoparasitoid used in biological control programs against medflies. Our results showed that *A. daci* progeny increased when parasitizing on CcaNV-infected larvae.

**Conclusions:** Our results proved that covert RNA viruses can impact on the insect ecology, directly affecting its insect host biology and indirectly influencing multitrophic interactions.

## Background

In the last decade, large-scale studies based on high-throughput sequencing data have redefined the virome of insects (Haoming et al., 2021; Käfer et al., 2019; Shi et al., 2016). The number of newly described insect-specific viruses (ISVs), especially those based on RNA (RNA-ISVs), has drastically increased (Haoming et al., 2021; Käfer et al., 2019; Shi et al., 2016). Despite the vast list of new virus species provided by these studies, very few studies have documented lethal or sublethal host effects associated with those viruses. Most RNA-ISVs are covert infections that do not cause noticeable symptoms to their hosts. Only in a few cases, RNA-ISVs have been found to induce behavioral and physiological changes in the host (Han et al., 2015). For instance, exposure to CO2 triggered the paralysis or death of various fruit fly species (Diptera: Tephritidae) infected with sigmavirus (*Rhabdoviridae*) (Longdon et al., 2012). In another example, the presence of the Kakugo virus (*Iflaviridae*) in the brain of the honey bee *Apis mellifera* L. (Hymenoptera: Apidae) has been correlated with the aggressive behavior of the bees (Fujiyuki et al., 2005).

In the agricultural pest *Ceratitis capitata* (Wiedemann) (Diptera: Tephritidae), also known as the Mediterranean fruit fly or medfly, up to 13 different RNA viruses have been described producing apparent covert infections without obvious symptoms in laboratory-adapted strains and field populations (Hernández-Pelegrín et al., 2022; Kondo et al., 2019; Llopis-Giménez et al., 2017; Sharpe et al., 2021). From these 13 RNA covert viruses described in medfly, only Ceratitis capitata nora virus (CcaNV) presence has been indirectly associated with a possible physiological cost for the host. A higher CcaNV abundance was reported in the group of medfly adults showing a shorter lifespan, while no correlation was found between CcaNV abundance and other parameters such as adult flight ability or mating behavior (Llopis-Giménez et al., 2017).

Viral infections in individuals of pest species may influence pest impact and population dynamics. For instance, the infection with Helicoverpa armigera iflavirus correlated with lower larval and pupal growth rates in the cotton bollworm *Helicoverpa armigera* (Hübner) (Lepidoptera: Noctuidae) (Yuan et al., 2020). In addition, viral infections may affect multitrophic interactions. A well-known example concerns the mutualistic viruses associated with parasitoid wasps (Volkoff & Cusson, 2020). Among those, dsDNA viruses are the best-studied examples since some parasitoid species require the presence of those viruses to complete their life cycle (Santos et al., 2022; Webb et al., 2006). These dsDNA viruses interfere with the host’s immune response and can be found integrated into the wasp genome (polydnaviruses) or as an exogenous virus (i.e., Diachasmimorpha longicaudata entomopoxvirus) (Coffman et al., 2020; Lawrence, 2002; Strand & Burke, 2020). Additionally, the presence of mutualistic RNA viruses of the *Reoviridae* family has been reported in the parasitoid *Diadromus pulchellus* (Wesmael) (Hymenoptera: Ichneumonidae) (Renault, 2012; Roossinck, 2011). Diadromus pulchellus reovirus 2 inhibits host-induced myelinization of the wasp egg, avoiding encapsulation and allowing further development of the parasitoid egg (Renault et al., 2005).

With or without the assistance of viruses, parasitic wasps can naturally reduce the populations of tephritid fruit flies. In this context, the release and conservation of natural parasitoids is a successful tool for tephritid pest management, including medfly (de Pedro et al., 2018; El-Heneidy et al., 2014; Garcia et al., 2020; Harbi et al., 2018; S. Ovruski et al., 2000; S. M. Ovruski & Schliserman, 2012; Vargas et al., 2012). So far, no viruses have been associated with the larval-pupal endoparasitoid *Aganaspis daci* (Weld) (Hymenoptera: Figitidae), which is used in biological control programs against medflies (de Pedro et al., 2018). In addition, it is unknown whether viruses present in medflies affect the interactions between medflies and the parasitoid wasp *A. daci*.

The release and conservation of fruit fly parasitoids combined with other pest control methods, such as the sterile insect technique (SIT), can contribute to a more robust pest management strategy (Argov & Gazit, 2008). SIT is the most widespread medfly control method based on the systematic area-wide release of sterile males obtained from mass-rearing facilities (Enkerlin, 2005). In insect mass-rearing facilities, the direct impact of covert viral infections on insect physiology may negatively affect mass production and/or alter the competitiveness of males released into the field for SIT applications. For instance, changes in insect rearing conditions can lead covert infections to trigger disease outbreaks and cause the collapse of the insect colonies (Bertola & Mutinelli, 2021; Maciel-Vergara & Ros, 2017).

In this study, we aim to assess the direct and indirect effects of a covert infection with an RNA virus on the host insect’s physiology and the interaction with another trophic level. We selected the CcaNV-medfly system as a model, purified CcaNV from a naturally infected medfly strain, and tested its transmission routes. After verifying the horizontal transmission of this virus, we orally infected CcaNV-free flies with CcaNV and studied the possible non-lethal effects caused by viral infection on host physiology. We also investigated the influence of the viral infection in medflies on the parasitism performance of the endoparasitoid *A. daci*. Overall, our results reported fitness costs associated with a CcaNV infection in medflies and furthermore revealed a role of this virus in multitrophic interactions potentially regulating medfly population dynamics in the field.

## Results

### Vertical and horizontal transmission of CcaNV

The presence of CcaNV and CcaIV2 in *C. capitata* eggs in the Vienna 8A strain, that naturally carries a covert infection with both viruses (Hernández-Pelegrín et al., 2022), was investigated by quantitative PCR (RT-qPCR). CcaIV2 was selected as positive viral control since previous works showed its ubiquitous distribution in medfly populations (Hernández-Pelegrín et al., 2022). According to our results both viruses were present in medfly eggs, with a higher relative viral abundance in the case of CcaIV2 (Figure 1A). To test whether viral vertical transmission occurred within the egg (transovarial) or via the eggshell (transovum), we evaluated the presence of the two viruses in dechorionated eggs. Both viruses (CcaNV and CcaIV2) were detected in chorionated as well as dechorionated eggs. Moreover, no significant differences in relative viral abundance were found between chorionated and dechorionated eggs, indicating the transovarial vertical transmission of these viruses (Figure 1A).

**Figure 1.**
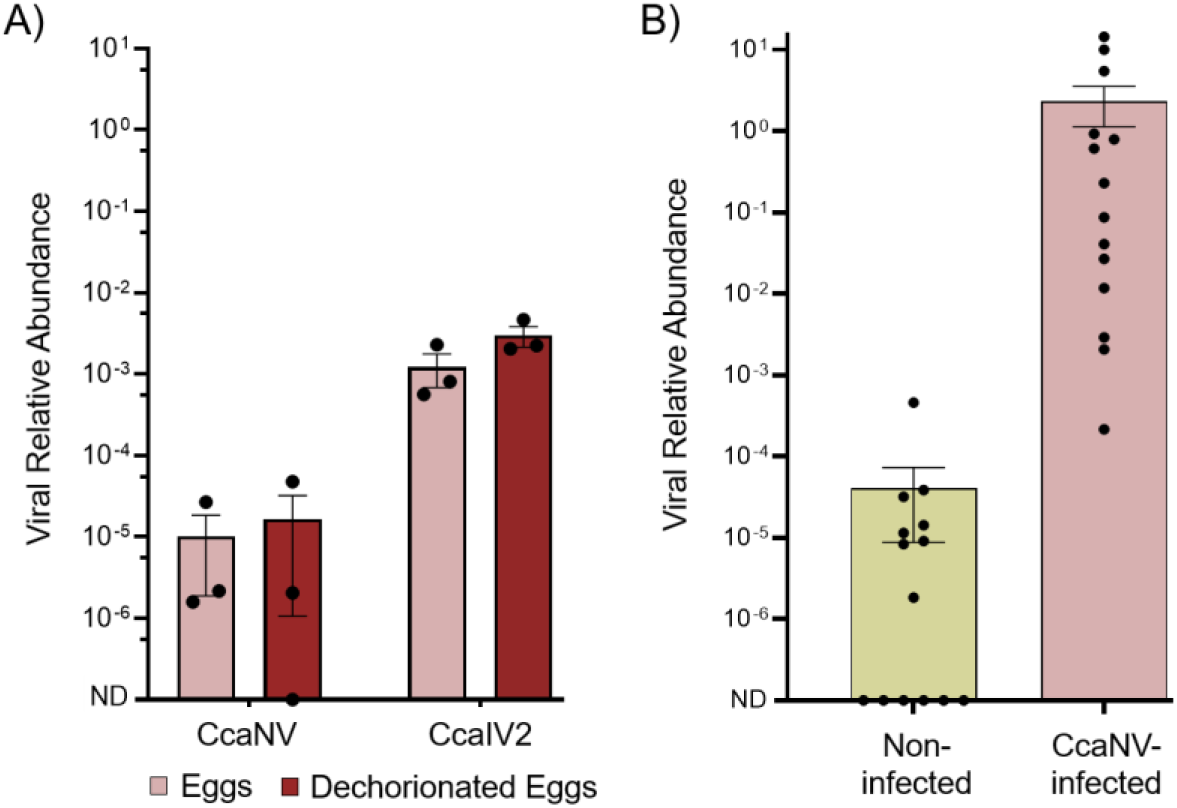
Analysis of vertical transmission of CcaNV and CcaIV2 and horizontal transmission of CcaNV. A) Viral relative abundance in *C. capitata* eggs (light red) and dechorionated eggs (dark red) of the Vienna 8A medfly strain. B) Viral relative abundance in pools of 5 pupae of the Control medfly strain before (non-infected) and after (CcaNV-infected) oral infection with a viral purification containing 10^9^ CcaNV genomes/μl mixed with the larval diet.

To analyze whether CcaNV can also be horizontally transmitted when medflies are feeding on a virus-infected diet, a viral purification containing CcaNV was obtained from the Vienna 8A medfly strain, mixed with the larval diet, and offered to first instar larvae from the CcaNV-free strain (named as Control strain). Larvae fed on the diet containing viral particles showed a dramatic increase in CcaNV titers at the pupal stage, demonstrating CcaNV horizontal transmission (Figure 1B).

### CcaNV infection has a direct impact on medfly survival

The influence of a CcaNV infection on different parameters related to medfly development (hatching rate, pupal weight, adult emergence) and medfly adults’ performance (flight ability and longevity under starvation stress) was analyzed (Figure 2). The hatching rate was significantly higher in CcaNV-infected flies (93.6% ± 0.7) when compared to the non-infected flies (89.8% ± 1.2) (*t* = 2.875; *df* = 14; *P* = 0.012). No delay or quickening in the developmental time from the egg to the pupal stage was observed for the CcaNV-infected group compared to the non-infected group. However, a significant difference was observed in terms of pupal weight (*t* = 8.029; *df = 14*; *P* < 0.001), with pupae deriving from CcaNV-infected larvae (9.79 ± 0.03 mg/pupae) being heavier than the non-infected pupae (9.26 ± 0.05 mg/pupae). No significant differences were observed in the percentage of emerged adults between CcaNV-infected (78.1 ± 6.8) and non-infected (92.1 ± 1.1) flies (*t* = 2.031; *df = 14*; *P* = 0.062) (Figure 2). Results of the flight ability, measured as the flight index (Parker et al., 2020), showed no significant differences (*t* = 1.271; *df= 10*; *P* = 0.233) between the CcaNV-infected (86.1 ± 3.4) and non-infected (89.6 ± 2.7) flies. In contrast, longevity under starvation stress was clearly affected by the presence of a CcaNV infection. For both CcaNV-infected and non-infected groups, flies died between day 3 and day 7 after adult emergence when exposed to starvation stress. However, considering day 4 post-emergence, when adult flies have reached their sexual maturity under natural conditions, up to 64% of the CcaNV-infected flies had died, compared to 43% of the non-infected flies (*t* = 3.745; *df = 30*; *P* = 0.008) (Figure 2).

**Figure 2.**
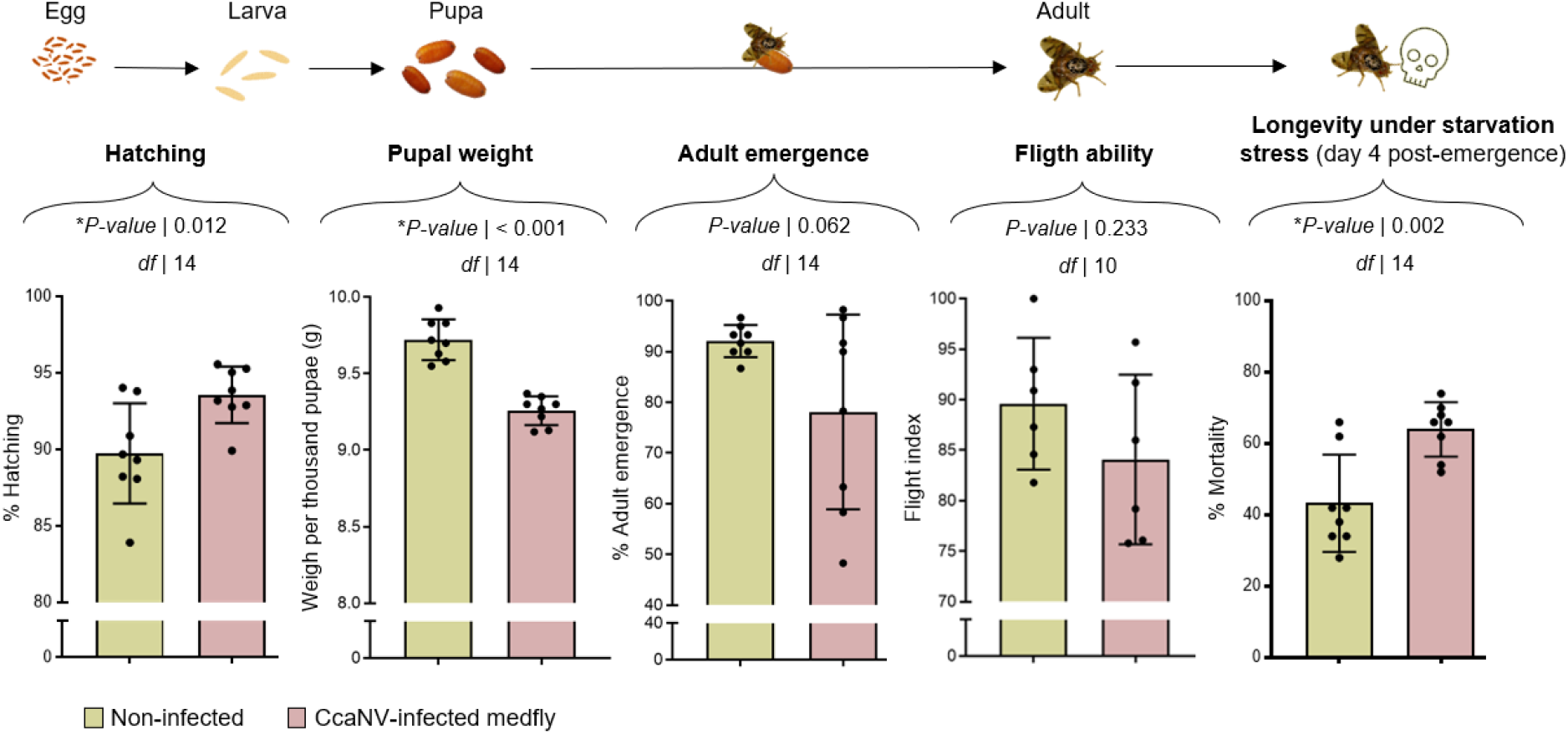
Influence of a CcaNV infection on the development and performance of medflies. The parameters analyzed were the hatching rate (number of hatched larvae/number of eggs), the pupal weight (g/1000 pupae), the adult emergence rate (number of emerged adults/number of pupae), the flight ability (flight index), and the longevity under starvation stress (mortality at day 4 post-emergence). Statistical differences between CcaNV-infected and non-infected groups were calculated using unpaired *t*-tests, and the degrees of freedom (*df*) and *P*-values are shown above each graph.

In summary, our results revealed that a covert infection with CcaNV alters certain biological parameters of the medfly host. We observed negative effects of the insect-virus interaction on host fitness, including a decreased pupal weight and shorter longevity under stress, but also a positive impact as seen for the increased hatching rate. To unravel the overall effect of a CcaNV infection, we estimated the probability of survival (PS) from eggs to the sexually active adult stage by jointly analyzing the hatching rate, adult emergence, and longevity (see Materials and Methods). Significant differences were observed between CcaNV-infected and non-infected groups after analyzing the 8 replicates performed per condition (*t* = 4.852; *df = 14; P* < 0.001). The PS estimated for CcaNV-infected flies was 0.26 ± 0.04, meaning that from an initial cohort of eggs, only 26% arrived until fertile adulthood in the above-mentioned conditions. On the other hand, the PS estimated for the non-infected flies was 0.47 ± 0.09.

### *CcaNV infection alters parasitism by* A. daci

To explore the indirect effects of a CcaNV infection on medfly ecology, we analyze CcaNV influence on the interaction between medflies and one of its main natural parasitoids, *A. daci*. The influence of a CcaNV infection on medfly attractiveness to *A. daci* females was assessed using an olfactometer assay. Results revealed that female *A. daci* adults were more attracted to the medfly diet containing CcaNV-infected larvae (n=22; 73%), in comparison to the diet with non-infected larvae (n = 8; 27%) (*χ*^2^ = 13,067; *df* = 1; *P* < 0.001) (Figure 3).

**Figure 3.**
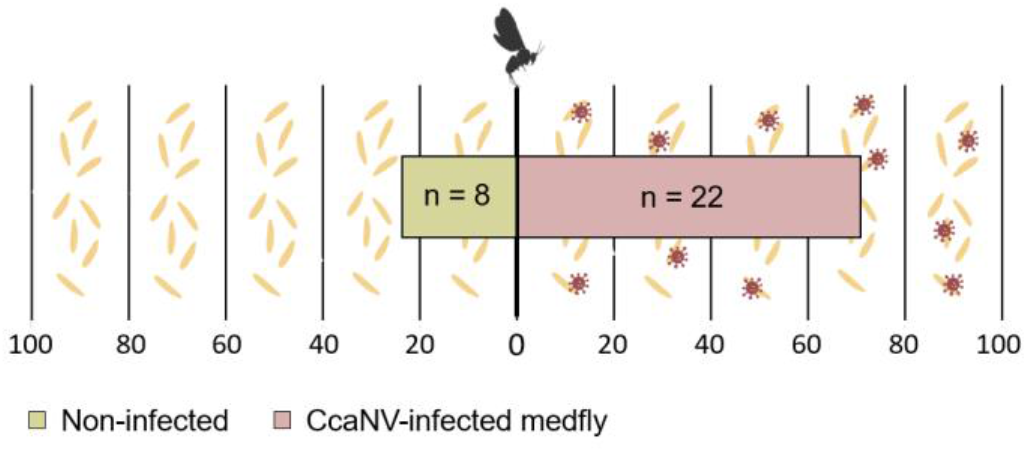
Response of *A. daci* female adults towards medfly diet containing non-infected or CcaNV-infected larvae of the Control strain, respectively. A total of 30 responses were measured.

After larvae recognition, parasitoids initiate oviposition into the larval bodies. We performed two different types of experiments to determine the influence of CcaNV infection in the parasitoid fecundity, defined as the number of parasitoid eggs laid by parasitoid females; and the parasitoid progeny, defined as the number of parasitoid adults able to emerge from medfly pupae. First, CcaNV-infected or non-infected larvae were offered separately to each male/female *A. daci* couple (non-choice assay). A first group of medfly larvae was used to determine parasitoid fecundity, and a second group was used to assess parasitoid progeny. Our results showed that the total number of oviposition scars per parasitoid female was significantly higher for pupae developed from CcaNV-infected larvae (64.51 ± 5.12) than from non-infected larvae (49.89 ± 3.22) (*F*_(1,61)_ = 10.05; *P* = 0.002). In accordance, CcaNV-infected larvae showed 2.75 ± 0.16 oviposition scars per pupa, significantly higher than the 2.41 ± 0.09 oviposition scars per pupa observed in the non-infected group (*F*_(1,61)_ = 5.77; *P* = 0.019) (Figure 4). Accordingly, significant differences were retrieved between groups in the percentage of wounded pupae (*F*_(1,61)_ = 5.76; *P* = 0.019), with 79.58 % ± 2.75 of the pupae presenting oviposition scars in the CcaNV-infected group and 70.33% ± 2.94 in the non-infected group (Figure 4).

**Figure 4.**
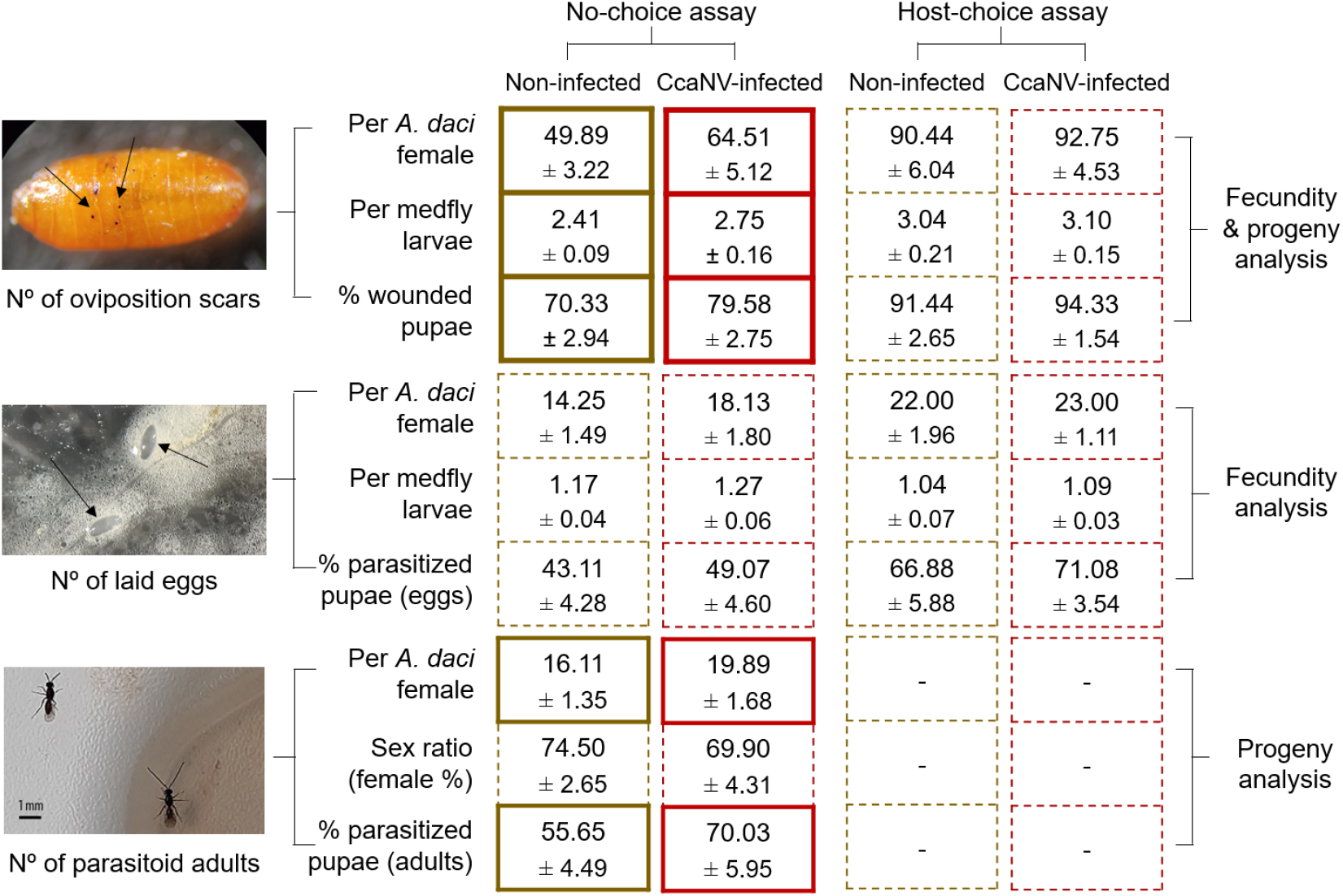
Influence of CcaNV infection on the parasitism of the parasitoid *A. daci*. Fecundity results refer to the number of eggs laid by parasitoid females, while progeny results focus on the new parasitoid adults emerging. Values are shown together with their standard error. Parameters presenting statistically significant differences between non-infected and CcaNV-infected groups are highlighted using continuous bolded lines in the table.

However, those differences in the number of oviposition scars were not reflected in the number of parasitoid eggs laid into the medfly larvae. Although there was a general tendency towards an increase in parasitoid fecundity on CcaNV-infected larvae, no significant differences were observed for the number of eggs laid by each *A. daci* female (*F*_(1,29)_ = 2.84; *P* = 0.102), the number of eggs found per pupa (*F*_(1,29)_ = 2.05; *P* = 0.163), or the percentage of parasitized pupae, defined as pupae containing at least one parasitoid egg (*F*_(1,29)_ = 0.97; *P* = 0.332) (Figure 4). In contrast, the progeny analysis that reflects the result of the whole parasitism process showed that both the number of emerged parasitoid adults per *A. daci* female (*F*_(1,32)_ = 7.66; *P* = 0.009) and the percentage of parasitized pupae giving rise to new *A. daci* individuals (*F*_(1,32)_ = 8.81; *P* = 0.005) were significantly different between groups. These parameters indicated an increase of parasitism success in medfly larvae infected with CcaNV since 70.03% ± 5.95 of the CcaNV-infected medfly larvae offered to *A. daci* were successfully parasitized in comparison to the 55.65% ± 4.49 of non-infected larvae. The sex ratio of the emerged parasitoid adults was biased towards females (74.50 ± 2.65% and 69.90 ± 4.31% for non-infected and CcaNV-infected groups, respectively) but was not linked to the presence of CcaNV infection in the parasitized medfly larvae (*F* _(1,32)_ = 0.82; *P* = 0.370) (Figure 4).

To complement the previous analyses, we explored the influence of CcaNV infection when an identical number of CcaNV-infected and non-infected medfly larvae were simultaneously offered to each male/female *A. daci* couple (host-choice assay). Compared to the results of the non-choice assay, the number of oviposition scars was higher in the host-choice assay in both CcaNV-infected and non-infected groups (F_(1,96)_ = 42.62; *P* < 0.001)). In contrast to the non-choice assay and probably caused by the high number of oviposition scars, the presence of CcaNV infection did not induce significant differences in *A. daci* fecundity for any of the selected parameters: number of oviposition scars by each parasitoid female (*F*_(1,29)_ = 0.10; *P* = 0.758), number of oviposition scars measured per medfly pupa (*F*_(1,29)_ = 0.07; *P* = 0.969), percentage of wounded pupae (*F*_(1,29)_ = 0.89; *P* = 0.352); number of eggs laid by each *A. daci* female (*F*_(1,29)_ = 0.21; *P* = 0.650), number of eggs per medfly pupa (*F*_(1,29)_ = 0.45; *P* = 0.508), and percentage of parasitized pupae containing at least one *A. daci* egg (*F*_(1,29)_ = 0.43; *P* = 0.518) (Figure 4).

## Discussion

Covert virus infections have been widely documented for different insect species, but little is known about their effect on host development and ecology. To further investigate the impact of covert virus infections on their host, we have studied the interaction between the agricultural pest *C. capitata* and Ceratitis capitata nora virus (CcaNV) and unravelled direct as well as indirect biological costs and benefits to the host associated with the covert infection with this virus.

Covert infections with certain viruses are always present in medflies, while others are present in some medfly strains but absent in others. For instance, CcaIV2 is present in all the strains and field populations analysed worldwide (Hernández-Pelegrín et al., 2022). In contrast CcaNV, can be either absent (Control strain) or present (Vienna 8A strain). In this study, a viral purification was obtained from the Vienna 8A strain and added to the Control strain’s larval diet, resulting in a significant increase of CcaNV in the Control strain, with no difference in the abundance of other RNA viruses such as the CcaIV2. A possible reason to explain this result is that, except for CcaNV, the RNA viruses presented in the viral purification were not able to orally infect larval stages of medfly. Alternatively, the concentration of these viruses in the purification may not have been high enough to increase the levels of the viruses already present on the Control strain. Further research would be needed to confirm the transmission route of the multiple viruses covertly infecting the medflies.

After confirming CcaNV horizontal transmission, we unravelled the effects caused by the infection in the medfly host. Although no lethal effects were associated with CcaNV, we observed a negative impact on the survival of the flies. First, CcaNV-infection caused a decrease in pupal weight, while previous studies highlighted the improved fitness of larger insects marked by a longer lifetime and a higher reproductive rate (Beukeboom, 2018). Similarly, a decrease in adult survival under starvation stress was observed in CcaNV-infected medflies. On the other hand, our analysis showed a positive effect of a CcaNV infection on the hatching rate, which was higher for CcaNV-infected larvae. However, the combined analysis of all measured parameters showed an approximate 50% reduction in the probability of survival of CcaNV-infected flies (PS = 47%) compared to non-infected flies (PS = 26%). Whether the reduced longevity observed in CcaNV-infected flies is caused by the decrease in weight due to the viral infection or the direct impact of the virus on adult survival under stress remains to be elucidated. In any case, these results suggest that a CcaNV infection could cause a competitive disadvantage to infected flies in the field, where the chances to face environmental stressors such as the absence of nutrients or the presence of natural parasitoids arise.

Until now, the study of mutualistic relationships between viruses and parasitoids had focused on parasitoid viruses. This is the case for polydnaviruses (PDVs), a group of dsDNA viruses that are integrated into the genome of parasitoid wasps and are required for successful parasitism of the parasitoid’s hosts (Strand & Burke, 2015). Apart from PDVs, exogenous viruses have also been associated with parasitoid wasps as in Diachasmimorpha longicaudata entomopoxvirus (DIEPV), an exogenous dsDNA virus from *Diachasmimorpha longicaudata* required for the successful parasitism of fruit flies (Coffman & Burke, 2020). Here, we focused for the first time on the influence of covert RNA viruses infecting the parasitized host on the performance of the parasitoid. In this context, we studied the interaction between medfly and *A. daci*, a parasitoid of fruit flies that effectively parasitize *C. capitata* (de Pedro et al., 2018). We observed a higher attraction of *A. daci* females towards CcaNV-infected medfly larvae in the olfactometer assay, which translated into a higher number of oviposition scars in the CcaNV-infected group when CcaNV-infected and non-infected larvae were offered separately (non-choice assays). In contrast, no differences in the number of oviposition scars were observed when CcaNV-infected and non-infected larvae were simultaneously provided to the parasitoid (host-choice assay). In this vein, a higher parasitism rate was observed in host-choice assays compared to non-choice assays, independently from the treatment. It is likely that the high rate of parasitizing attempts (high number of scars) was buffering the effect of the CcaNV infection on the host preference. Given the reduced space of the experimental arena and the high density of host larvae, it is possible that *A. daci* females were unable to distinguish the source of the attractant volatiles when CcaNV-infected and non-infected larvae were simultaneously offered. Further research on the blend of volatiles emitted by CcaNV-infected larvae would help to understand this response. Alternatively, *A. daci* females may have prioritized oviposition in CcaNV-infected larvae, and, afterward, oviposit in larvae of the non-infected group.

During non-choice assays, the higher attraction and number of oviposition scars observed in CcaNV-infected larvae did not correlate with a higher number of laid eggs but correlated with a higher emergence of parasitoid adults. Considering these results, it is tempting to hypothesize that *A. daci* development is favoured inside the infected larvae since a higher number of parasitoids emerged in the CcaNV-infected group, while no significant differences were displayed in the number of laid eggs. This could result from a weaker immune response activation against the parasitoid motivated by the presence of the covert infection. The analysis of the expression levels of medfly immune system markers after parasitism would help to confirm this hypothesis. However, while the differences observed in the number of laid eggs were not statistically significant, the values for the different fecundity parameters were higher in the CcaNV group. In this scenario, the higher attraction towards infected flies could led to more oviposition scars, laid eggs, and parasitoid progeny.

In any event, our results uncovered CcaNV as the first instance of a covert RNA virus benefiting the parasitism of its host. Whether these results can be extrapolated to other parasitoid-host systems will need further confirmation. In this context, a further investigation of the interactions between covert viruses present in the host and the symbiotic viruses present in the parasitoid will be of great interest. The presence of covert RNA viruses is extensive in tephritid fruit flies (Haoming et al., 2021; Sharpe et al., 2021; Zhang et al., 2022) and may influence the relations between parasitoids and fruit fly hosts in the field. Additionally, we must recognize the possibility that the observed positive effect of CcaNV on *A. daci* parasitism performance depends on the environmental context. For instance, a PDV infection in the white butterfly parasitoid *Cortesia glomerata* (L.) (Hymenoptera: Braconidae) positively contributes to *C. glomerata* parasitism but, in a multitrophic level, exerts a negative effect in offspring survival by induction of plant volatiles and increase attraction to the *C. glomerata* hyperparasitoid wasp, *Lysibia nana* (Gravenhorst) (Hymenoptera: Ichneumonidae) (Zhu et al., 2018).

Our results have revealed that a covert infection with CcaNV negatively impacts medfly survival and contributes to increased parasitism by one of its parasitoids, *A. daci*. Previous studies on the distribution of CcaNV confirmed its presence in different medfly strains, including flies collected from the field and flies mass-reared for SIT (Hernández-Pelegrín et al., 2022). It is expected that covert CcaNV infections in field individuals would negatively impact the field populations and influence medfly ecology. From a pest control point of view, our results suggest that CcaNV infections in the field, as well as the continuous inoculation of CcaNV in the field populations through the release of infected males for SIT, would contribute to successful biological control of the pest. In contrast, a CcaNV infection would also reduce longevity under starvation stress of the infected males released for SIT. The trade-off between both aspects would define whether the presence of a covert infection with CcaNV is advantageous or not for area-wide management using SIT. Alternatively, we cannot discard that this negative effect caused by CcaNV to the flies is surpassed with a positive impact that has still to be unraveled.

To sum up, the direct and indirect effects observed after infection with CcaNV highlight the impact of covert RNA viruses on medfly ecology and set the basis for a further and necessary characterization of covert virus-host interactions.

## Conclusions

In this study, we moved beyond viral detection to present a comprehensive study of the multitrophic effects associated with a Ceratitis capitata nora virus (CcaNV) covert infection in the medfly. Despite the absence of lethal effects, the specific reduction in pupal weight and survival under starvation stress reported for CcaNV-infected flies resulted in a significantly lower probability of survival. Moreover, the CcaNV-infection enhanced the attractiveness of medfly larvae towards its parasitoid *A. daci*, although the number of eggs laid by the parasitoid was not significantly different. Instead, a higher number of parasitoid adults emerged from CcaNV-infected larvae suggesting that CcaNV favours the endoparasitoid development inside the medfly larvae. In conclusion, this work proves the relevance of covert viral infections on the modulation of host development and multitrophic interactions. Further research, including different insect-virus systems, will contribute to better understand the biological role of covert viral infections in insects.

## Experimental procedures

### 2.1 Ceratitis capitata *insects*

Two different medfly strains (Control and Vienna 8A) were selected for the experiments based on the presence or absence of CcaNV infection in these strains. The Control strain was chosen as a CcaNV-free strain based on the results of viral abundance analysis. This strain was subsequently infected with purified CcaNV (see below) to perform fitness and parasitism experiments. The Control strain originates from flies collected from experimental fields at the “Instituto Valenciano de Investigaciones Agrarias” (IVIA) in 2001, and it has been reared since then under laboratory rearing conditions of 26°C, 40 to 60% humidity, and 14/10h light/dark cycles (Arouri et al., 2015). The colony is regularly renewed with field-captured flies from the experimental fields at the IVIA. The Vienna 8A (V8A) strain was selected for the analysis of vertical transmission based on its relevance for the application of SIT program for the control of *C. capitata* in the Valencian community. Regarding its origin, V8A is a naturalized V8 strain derived from mixing the temperature-lethal genetic sexing strain Vienna-8 mix 2002 with wild individuals collected in citrus orchards in the province of Valencia, Spain. This strain is produced at the mass-rearing facility located in Caudete de las Fuentes (Valencia, Spain), which is financed by the Department of Agriculture of Valencia and implemented by the state-owned company Empresa de Transformación Agraria S.A. (Grupo TRAGSA, Valencia, Spain) (Pla et al., 2021). Eggs from the Vienna 8A population were received from the mass-rearing facility and reared under the above-mentioned laboratory rearing conditions.

### 2.2 Aganaspis daci *insects*

The selected *A. daci* strain originates from medfly larvae collected in figs from the nearby area of Bétera (Valencia, Spain) in 2010. It is maintained under laboratory-rearing conditions (27 ± 2 °C, 65% ± 10% RH, 16:8 (L:D) photoperiod) in the IVIA facilities, using the IVIA medfly strain as host and fed with a mixture of wheat bran, sugar, and brewer’s yeast (Noriega, 2017).

### 2.3 CcaNV detection and quantification in medfly samples

The presence of CcaNV in medfly was determined using molecular methods as previously described in Hernández-Pelegrín et al., 2022. Briefly, total RNA was isolated from individual larvae or pupae using TriPure isolation reagent (cat. no. 11667157001; Roche, Mannheim, Germany) according to the manufacturer’s protocol. DNAse treated-RNA was reverse transcribed into cDNA using random hexamers and oligo (dT) primers with the Prime-Script RT Reagent Kit (Perfect Real Time from Takara Bio Inc., Otsu Shiga, Japan). Viral presence was assessed through RT-qPCR (StepOnePlus Real-Time, Applied Biosystems, Foster City, CA) by preparing 20μL reactions containing 5x HOT FIREpol EvaGreen qPCR Mix Plus (ROX) from Solis BioDyne (Tartu, Estonia), 2μL of primers and 4μL of cDNA. Specific primers were used to amplify the coding region of the RNA-dependent RNA polymerase of CcaNV and CcaIV2 (Llopis-Giménez et al., 2017). The medfly ribosomal L23a gene was amplified as an endogenous control of the RNA concentration using available primers (Llopis-Giménez et al., 2017). The relative viral abundance was calculated by comparison of viral Ct values and L23a Ct values, according to the 2^-ΔCT^ method described by Livak and Schmittgen (Livak & Schmittgen, 2001). Data were visualized and analysed using GraphPad Prism version 8.0.0 for Windows (GraphPad Software, San Diego, California USA, www.graphpad.com).

### 2.4 CcaNVpurification

A viral purification containing CcaNV was obtained from pools of fifty pupae from the Vienna 8A strain, previously stored at 20°C, following the protocol described in Hernández-Pelegrín et al., 2022. After purification, the viral pellets were resuspended in 2ml of PBS and stored at −80°C until use. Aliquots of 100 μL were used for RNA extraction and virus quantification through RT-qPCR as described above. Standard curves were obtained to quantify the viral genomes per μg of RNA in the viral purifications by using a fixed number of viral genome copies cloned in the pGEMTeasy vector (Llopis-Giménez et al., 2017).

### 2.5 Eggs dechorionation

To distinguish between transovarial and transovum transmission routes, viral presence was assessed in dechorionated eggs of the Vienna 8A strain. To do that, eggs were collected and submerged in 50% sodium hypochlorite for 2 minutes to eliminate the chorion. Then, eggs were washed 3 times in milli-Q water during 3 min, and once with phosphate buffered saline (PBS). During the treatments, the eggs were maintained in a shaking platform to favour the exposure of the eggs to the solutions. Same protocol was performed for non-treated eggs substituting the 50% sodium hypochlorite for milli-Q water.

### 2.6 Oral infection of medfly larvae with CcaNV purification

Medfly artificial diet was prepared with a combination of nutritional elements (wheat bran, sugar, and brewer’s yeast); and preservative and antiseptic compounds (methylparaben, sodium propylparaben and sodium benzoate) (de Pedro et al., 2017). For each bioassay, the same batch of artificial diet was separated into two stocks: non-infected and CcaNV-infected. For each assay, 100g diet was inoculated with 1 ml of the viral stock at 10^9^ CcaNV genomes/μL, diluted with 3ml of water, and thoroughly mixed with the artificial diet using a flask and a glass rod. To maintain moisture conditions in the controls, 4ml of water were mixed with 100g of the non-infected diet stock. For each assay, the 100g of diet were divided into two technical replicates by placing 50g of diet in 9cm petri dishes. About thousand fresh eggs from the *C. capitata* Control strain were collected, mixed with 2 ml of water, and added to one replicate per condition. 24h later, the process was repeated with newly collected eggs and the remaining replicate. CcaNV-infected and non-infected groups were maintained in separate incubators under the same rearing conditions (26°C, 40 to 60% humidity, 14/10h light/dark cycles) to prevent viral contaminations. Larvae of each batch were collected for confirmation of the CcaNV-infection. CcaNV infection was confirmed in pools of 5 larvae, with two replicates per batch, using RT-qPCR as mentioned above.

### 2.7 Medfly fitness analysis

Fitness parameters were selected based on their potential implications in medfly ecology, the regular operation of mass-rearing facilities, and the field application of SIT-based control (FAO/IAEA/USDA, 2019). These parameters were subjected to a comparative analysis between CcaNV-infected and non-infected flies to determine the influence of viral infection on normal medfly development. Both experimental groups were always handled side-by-side, and flies were laboratory-reared under controlled conditions. For all the fitness parameters under analysis, the statistical differences between the groups were tested with GraphPad Prism version 8.0.0 for Windows (GraphPad Software, San Diego, California USA, www.graphpad.com) using an unpaired t-test (Parker et al., 2020).

#### 2.7.1 Hatching rate

Random cohorts of approximately 150 eggs deposited by CcaNV-infected and non-infected medflies, respectively, were placed in 9 cm petri dishes covered with dark and wet filter paper at the bottom. The experiment was revised daily for five days to annotate the number of hatched larvae and calculate the hatching rates. Two independent biological replicates were performed consisting of 4 cohorts of 150 eggs per condition and replicate.

#### 2.7.2 Pupal weight

Cohorts of sixty pupae were weighted 5 days after the first larvae jump, which is the process before pupa formation. Four cohorts per treatment were used, and the experiment was repeated twice. The total weight of the cohorts divided by the number of pupae provided the mean pupal weight per condition.

#### 2.7.3 Adult emergence

Pupae were collected for each treatment five days after the first larval jump. Then, pupae were incubated in 15cm petri dishes covered with muslin until adulthood or dead. Pupae development was checked daily, and the number of new adults was annotated to calculate the emergence percentages and the required time to complete the adult emergence. Two independent biological replicates were performed consisting of 4 cohorts of 60 pupae per condition and replicate.

#### 2.7.4 Flight ability

For the flight ability test, a petri dish containing 60 pupae was placed at the base of a black cylinder (9 cm in diameter, 10 cm high, and 3 mm thick walls) lightly coated with talcum powder on the inside walls to prevent flies from walking out the tube. The cylinder was introduced in a ventilated plexiglass cage (30 x 40 x 30 cm) under controlled conditions of 25 ± 2°C and 70% ± 10% relative humidity. For a period of 5 days from the first adult emergence, the medfly adults were counted daily in the initial tube, the plexiglass cage, and one additional tube aimed to identify re-entries (FAO/IAEA/USDA, 2019). Flight capacity was calculated using the following formula: (F + 2 x RE)/T, where F is the number of flies that flew out of the initial tube, RE is the number of flies that re-entered the new tube and T is the total number of flies including the flies which remained in the initial tube and the pupae that failed to complete their development. Two independent biological replicates consisting of three cylinders per condition and replicate were performed.

#### 2.7.5 Adult longevity under stress

Longevity was assessed for four groups of 25 males and 25 females per condition. Adult flies were maintained under the above-described rearing conditions, but water, sugar and protein were only provided for two days after emergence to generate a stress for the flies. Mortality was annotated at day 4 post-emergence. Two independent biological replicates were performed.

#### 2.7.6 Probability of survival (PS)

To study the overall effect of CcaNV infection, we defined the probability of survival from eggs to sexually active adult stage (PS) as % Hatching/100 x % Adult emergence/100 x (1 - % Mortality/100). From all the parameters under study, only those directly related with medfly survival were considered. PS was calculated for the eight replicates per condition.

### 2.8 A. daci *olfactory testing*

The response of *A. daci* to the olfactory stimuli triggered by viral infection was assessed with a Y-tube olfactometer (Analytical Research Systems, ARS It, Gainesville, FL, USA). The glass Y-tube was 2.4 cm in diameter, with two arms of 5,75 cm long and a 13.5 cm base. A unidirectional airflow of 150 ml/min was produced from the arms to the base using an air pump connected to two 5 liters crystal jars containing the odour sources (de Pedro et al., 2021). All the assays were performed under controlled conditions of 23 ± 2°C, 60% ± 10% relative humidity, and 2516 lux.

Forty-eight hours before the experiment, one hundred 3^rd^ instar medfly larvae from Control strain were placed in 9 cm petri dishes containing a) diet mixed with viral purification (CcaNV-infected) or b) diet mixed with water as control (non-infected). The diets were prepared as explained in the previous section, and each of them was introduced in one of the 5L crystal jars as odour source.

Regarding the parasitoid, 8-day-old *A. daci* females with no previous parasitic history on medfly larvae were collected for the experiment. Parasitoid females were individually placed in the base of the olfactometer, and their behaviour was observed for as long as 15 minutes. When the parasitoid individuals stayed for more than 10 seconds at 3 cm or longer from the bifurcation of the Y-tube, the response was considered positive. The experiment finished when 30 positive responses, meaning the parasitoid female selected one of the offering sides, were obtained. To minimize the effect of the experimental design on the parasitoid response, the connexion between the crystal jars containing the odour source and the Y-tube was inverted every five responses. Additionally, the connexion and the jars were cleaned with acetone every 10 attempts to eliminate the potential concentration of odours.

### 2.9 A. daci *parasitism assays*

To analyze the differences in *A. daci* parasitism performance between non-infected and CcaNV-infected groups, we designed parasitism units consisting of a plastic box with a muslin window on the upper surface (14,5 cm x 14,5 cm x 7,5 cm) (Figure S1, S2). Water and sugar were provided *ad libitum* inside the parasitism unit, and controlled conditions of 25 ± 2°C, 70% ± 10% relative humidity and 16:8 h (L:D) photoperiod were maintained. Two distinct assays were designed to understand the parasitism performance over non-infected and CcaNV-infected larvae offered simultaneously (host-choice assay) or separately (no-choice assay). For the host-choice assay, fifteen CcaNV-infected and fifteen non-infected larvae were simultaneously offered to a couple of *A. daci* adults (male and female) in each parasitism unit (Figure S1). The parasitoid adults selected were 6-8 days of age and had no previous contact with medfly larvae. After 24h, the 30 medfly larvae were removed, and new 3^rd^ instar larvae were offered to the same parasitoid couple for an additional 24h. Offered larvae were placed in two closed 5.5 cm petri dishes with two apertures covered with muslin, one per group. For the non-choice assay, fifteen medfly 3^rd^ instar larvae were mixed with larval diet and offered separately to one couple of *A. daci* adults each, in different parasitism units (Figure S2). For both assays, the larvae recovered after parasitism from each parasitism unit and treatment were incubated until pupation in ventilated petri dishes. Eight parasitism units were designed per each group of larvae (non-infected and CcaNV-infected). Two biological replicates at different time points (blocks) were performed for the host-choice assays and the no-choice assay for fecundity analysis. Instead, three blocks were prepared for the no-choice assays to study parasitoid progeny. As control for the natural mortality of the medfly larvae, two parasitism units containing no parasitoids were added to each assay. For the statistical analysis, the variability associated to the CcaNV-infection was considered as a fix factor while the variability associated to the different blocks was considered as a random factor and analyzed using a univariate mixed-model ANOVA test (Table S1).

#### 2.9.1 Fecundity

Fecundity assays were designed to determine whether *A. daci* oviposition is enhanced when medfly larvae infected with CcaNV were offered. The medfly larvae collected from the parasitism units were incubated until pupation. Five days after pupae formation, the oviposition scars produced by parasitoid females on the surface of the pupae were quantified with the help of a binocular (Leica M165C) to determine the number of oviposition scars. Then, medfly pupae were dissected using entomological tweezers to assess the number of *A. daci* eggs laid in each medfly pupa. On behalf of this data, we analysed the number of oviposition scars and parasitoid eggs per *A. daci* female, per medfly pupa, and the percentage of wounded pupa (presenting one or more oviposition scars) and parasitized pupa (containing one or more eggs of *A. daci*). The statistical differences between groups (non-infected and CcaNV-infected) were determined using a univariate mixed-model ANOVA test (Figure 4) and were considered statistically significant below the *P* < 0.05 threshold.

#### 2.9.2 Progeny

Progeny assays aimed to explore whether the emergence of *A. daci* adults varied due to CcaNV infection in medfly. First, the medfly larvae recovered from the parasitism units were incubated until pupation, and the oviposition scars on the surface of the pupae were counted using a binocular. Differently from fecundity analysis, pupae were incubated until completion of their development, which occurred about seven days later in the case of medfly emergence or 17-19 days later for the emergence of parasitoids. After 5 weeks, the number of adult parasitoids or medflies was counted, and the pupae which remained closed were dissected. Parasitoid adults completely formed inside the medfly pupae were considered successful parasitism since the emergence times for *A. daci* have a wide range. On behalf of the data recovered in this assay, we assessed the number of parasitoid adults that emerged per *A. daci* female, the sexual ratio of the *A. daci* adults that emerged and the parasitism efficiency, defined as the percentage of pupae leading to a successful parasitoid emergence. The statistical differences between groups (non-infected and CcaNV-infected) were determined using univariate mixed-model ANOVA test (Figure 4) and were considered statistically significant below the *P* < 0.05 threshold.

## Abbreviations

CcaIV2: Ceratitis capitata Iflavirus 2
CcaNV: Ceratitis capitata nora virus
ISVs: insect-specific viruses
IVIA: Instituto Valenciano de Investigaciones Agrarias
PDVs: Polydnaviruses
PS: the probability of survival from medfly eggs to the sexually active adult stage
SIT: Sterile Insect Technique

## Acknowledgments

We acknowledge Jaime García de Oteyza from TRAGSA for providing samples from their medfly mass-rearing facilities. We acknowledge Joop van Loon from the Wageningen University for his comments on the interpretation of results.

## Funding

This study was supported by the INSECT DOCTORS program, funded under the European Union Horizon 2020 Framework Program for Research and Innovation (Marie Sklodowska-Curie Grant agreement 859850) and projects IVIA-52202 and IVIA-51916 from Institute Valenciano de Investigaciones Agrarias (both projects are susceptible of being co-financed by the European Union through the FEDER). V.I.D.R. is supported by a VIDI-grant of the Dutch Research Council (NWO; VI. Vidi. 192.041).

## Author contributions

S.H., F.B., and L.H.-P. performed conceptualization of the research; the medfly strains used were maintained by F.B., M.C-O, and O.D; the methodology was designed and applied by: L.H-P., R.G-M., E.LL., L.N, Á.L.-G., and M.C-O; the original-draft was written by L.H-P., and reviewed and edited by S.H., V.I.D.R., F.B, M.C-O., A.U, and M.P-H. All the authors have read and agreed to the published version of the manuscript.

## Corresponding authors

Correspondence to Francisco Beitia or Salvador Herrero.

## Ethical declarations

Ethics approval and consent to participate: Not applicable.

Consent for publication: Not applicable.

Competing interests: The authors declare that they have no competing interests.

## Supplementary Information

*Table S1:* Estimation of the residual variability between biological replicates (blocks) for the parameters under analysis in the “non-choice” (a) and “host-choice” (b) fecundity analysis and the non-choice progeny analysis (c).

*Figura S1*. Parasitism unit for “host-choice” assay. A) Whole overview of the parasitism unit. B) inner view of the parasitism unit where both non-infected (larvae -) and CcaNV-infected larvae (larvae +) of control strain were simultaneously offered to a pair of *A. daci* parasitoids.

*Figura S2*. Parasitism unit for “non-choice” assay. A) Whole overview of the parasitism unit. B) Adapted opening where medfly larvae of control strain are offered to *A. daci* parasitoids. Each parasitism unit will contain larvae of one of the conditions: non-infected or CcaNV-infected.

## Notes

### Competing Interest Statement

The authors have declared no competing interest.

