## Supplementary Table 1, Supplementary Figures 1, 2 for "Beyond viral detection: multitrophic effects of covert infection with an RNA virus in medfly"

*Table S1*. Estimation of the residual variability between biological replicates at different time points (blocks) for the parameters under analysis in the "non-choice" (a) and "host-choice" (b) fecundity analysis and the non-choice progeny analysis (c).

| **A)** Variable | $\sigma^2$ Residual | $\sigma^2$ Block | Relative variability (%) |
|---|---|---|---|
| Nº of oviposition scars per *A. daci* female | 365,95 | 0,0025 | < 0,001 |
| Nº of oviposition scars per medfly pupa | 0,449 | 0,0169 | 0,052 |
| Percentage of oviposition scars per pupae | 338,19 | 0,0841 | 0,025 |
| Nº of eggs laid per *A. daci* female | 42,25 | 0,0625 | 0,148 |
| Nº of eggs laid per medfly pupa | 0,04 | < 0,001 | 2,50 |
| Percentage of parasitized pupa | 292,75 | 0,1521 | 0,052 |

| **B)** Variable | $\sigma^2$ Residual | $\sigma^2$ Block | **Relative variability** (%) |
|---|---|---|---|
| Nº of oviposition scars per *A. daci* female | 444,79 | 0,0480 | 0,010 |
| Nº of oviposition scars per medfly pupa | 0,518 | 0,040 | 7,722 |
| Percentage of oviposition scars per pupae | 75,17 | 0,0 | 0,0 |
| Nº of eggs laid per *A. daci* female | 38,19 | 0,122 | 0,319 |
| Nº of eggs laid per medfly pupa | 0,05 | < 0,001 | 2,0 |
| Percentage of parasitized pupa | 329,78 | 0,260 | 0,078 |

| **C)** Variable | $\sigma^2$ Residual | $\sigma^2$ Block | **Relative variability** (%) |
|---|---|---|---|
| Nº of oviposition scars per *A. daci* female | 350,81 | 1,69 | 0,479 |
| Nº of oviposition scars per medfly pupa | 0,260 | < 0,001 | < 0,001 |
| Percentage of oviposition scars per pupae | 105,062 | 1,060 | 0,998 |
| Percentage of parasitized pupa | 187,96 | 2,28 | 0,63 |
| Nº of adults per *A. daci* female | 13,987 | 2,689 | 16,120 |
| Sex ratio (% of females) | 230,43 | 0 | 0 |

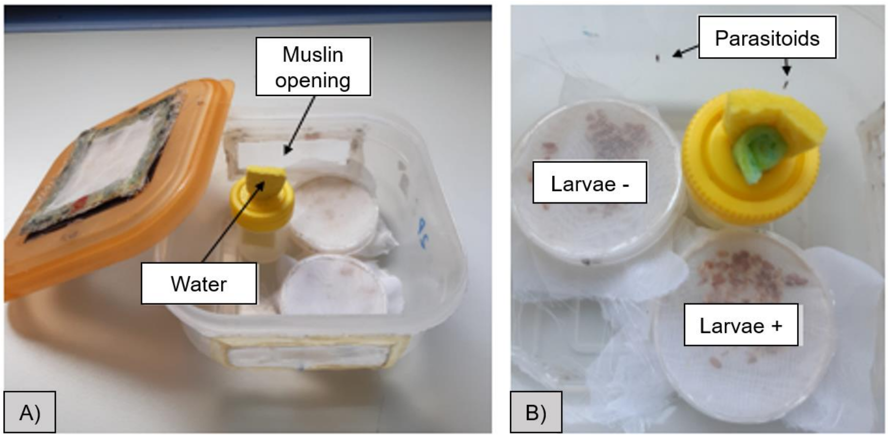


*Figura S1*. Parasitism unit for “host-choice” assay. A) Whole overview of the parasitism unit. B) inner view of the parasitism unit where both non-infected (larvae -) and CcaNV-infected larvae (larvae +) of control strain were simultaneously offered to a pair of *A. daci* parasitoids.

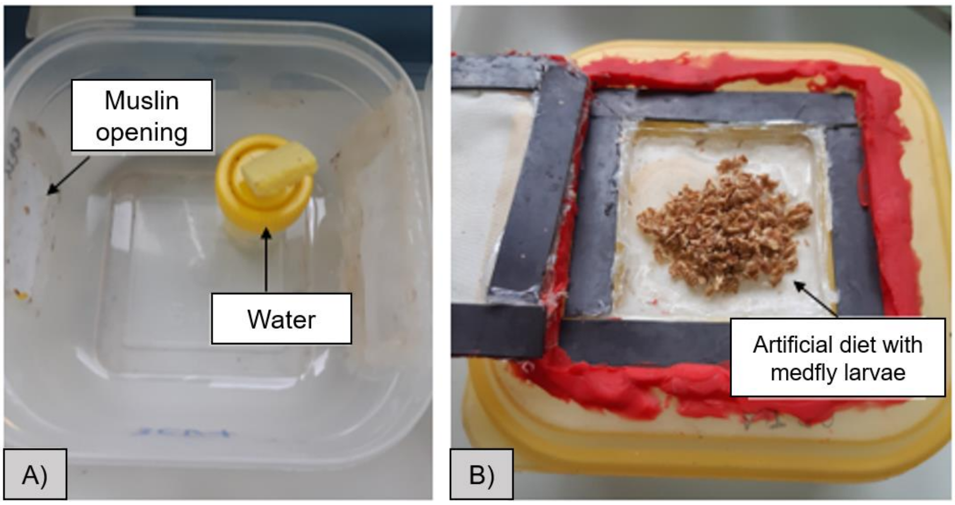


*Figura S2*. Parasitism unit for “non-choice” assay. A) Whole overview of the parasitism unit. B) Adapted opening where medfly larvae of control strain are offered to *A. daci* parasitoids. Each parasitism unit will contain larvae of one of the conditions: non-infected or CcaNV-infected.
